# Individual and combined effects of Cannabidiol (CBD) and Δ9-tetrahydrocannabinol (THC) on striato-cortical connectivity in the human brain

**DOI:** 10.1101/2020.11.20.391805

**Authors:** Matthew B. Wall, Tom P. Freeman, Chandni Hindocha, Lysia Demetriou, Natalie Ertl, Abigail M. Freeman, Augustus PM Jones, Will Lawn, Rebecca Pope, Claire Mokrysz, Daniel Solomons, Ben Statton, Hannah R. Walker, Yumeya Yamamori, Zixu Yang, Jocelyn L.L. Yim, David J. Nutt, Oliver D. Howes, H. Valerie Curran, Michael Bloomfield

**Author notes:** **Corresponding Author:** Matthew Wall, Invicro London, Burlington Danes Building, Hammersmith Hospital, Du Cane Road, London, W12 0NN, United Kingdom.

## Abstract

Cannabidiol (CBD) and Δ9-tetrahydrocannabinol (THC) are two major constituents of cannabis with contrasting mechanisms of action. THC is the major psychoactive, addiction-promoting, and psychotomimetic compound, while CBD may have somewhat opposite effects. The brain effects of these drugs alone and in combination are poorly understood. In particular the striatum is implicated in the pathophysiology of several psychiatric disorders, but it is unclear how THC and CBD influence striato-cortical connectivity. Across two placebo-controlled, double-blind studies, we examine the effects of THC, CBD, and THC+CBD on the functional connectivity of striatal sub-divisions (associative, limbic, and sensorimotor) using resting-state functional Magnetic Resonance Imaging (fMRI) and seed-based functional connectivity analyses. Study 1 (N=17; inhaled 8mg THC, 8mg THC+10mg CBD, placebo) showed strong disruptive effects of both THC and THC+CBD conditions on connectivity in the associative and sensorimotor networks, but a specific effect of THC in the limbic striatum, which was alleviated in the THC+CBD condition such that it did not differ from placebo. In Study 2 (N=23, oral 600mg CBD, placebo) CBD increased connectivity in the associative network, but relatively minor decreases/disruptions were found in the limbic and sensorimotor. In conclusion, THC strongly disrupts striato-cortical networks, and this effect is selectively mitigated in the limbic striatum when co-administered with CBD. When administered alone, 600mg oral CBD has a more complex effect profile of relative increases and decreases in connectivity. The insula emerges as a key region affected by cannabinoid-induced changes in functional connectivity, with potential implications for understanding cannabis related disorders, and the development of cannabinoid therapeutics.

## Introduction

Cannabis is a widely used recreational drug and has been used as such by humans for thousands of years for recreational, spiritual and medical purposes. The pharmacology of cannabis is complex, with almost 150 known cannabinoid compounds present in naturally occurring cannabis plant matter (Hanuš et al., 2016). Two of the major naturally occurring cannabinoids are Δ^9^-tetrahydrocannabinol (THC) and cannabidiol (CBD). THC is the major psychoactive compound and is responsible for the majority of the subjective and cognitive effects (Curran et al., 2002), including apathy, feeling ‘stoned’, amnesia, anxiety, and psychotomimetic effects (D’Souza et al., 2004). THC is thought to exert its effects primarily by partial agonism at the CB1 receptor (Pertwee, 2008). CBD has less well understood and more complex pharmacological effects, including negative allosteric modulation at the CB1 receptor (Chesney et al., 2020), reducing reuptake of anandamide, and action on GPR55, μ-opioid, and 5-HT1A receptors (Pertwee, 2008). CBD has antipsychotic, (Leweke et al., 2012; McGuire et al., 2018), anxiolytic (Bergamaschi et al., 2011a) and anti-addictive (Hindocha et al., 2018; Hurd et al., 2019; Freeman et al., 2020) properties, and therefore has broadly oppositional neuropsychopharmacological effects to THC (Curran et al., 2016; Gunasekera et al., 2020). Experimental studies co-administering THC and CBD have produced mixed results, but the most common finding was that CBD reduced the effects of THC (Freeman et al., 2019b).

Cannabis is currently moving towards a decriminalised or fully legal status in a number of jurisdictions. There is also renewed interest in the medical uses of cannabinoids, with growth in their medical licensing (Hasin et al., 2017; Lucas & Walsh, 2017; Freeman et al., 2019a), particularly for the treatment of chronic and neuropathic pain (Leung, 2011) and mental health conditions (Walsh et al., 2017). As use of cannabinoids in medical contexts becomes more widespread, it is vital to understand the intricate pharmacological and physiological mechanisms behind their potential therapeutic effects. One brain system known to be strongly affected by both acute and chronic use of cannabis of particular relevance to therapeutic, recreational, and harmful effects is the dopaminergic system and associated brain regions, principally the striatum (Bloomfield et al., 2018). The density of CB1 receptors is medium to high in striatal regions (Glass, Dragunow & Faull, 1997) and previous work has shown reductions in striatal dopamine function in cannabis users (Bloomfield et al., 2014; Tomasi, Wang & Volkow, 2015; Van De Giessen et al., 2017), and selective dopamine release in the limbic subdivision of the striatum with an acute THC challenge (Bossong et al., 2015). Functional and behavioural data have also shown that cannabis can acutely modulate striatal responses to hedonic stimuli (Freeman et al., 2017), and impair reward learning (Lawn et al., 2016). Multiple lines of evidence implicate the striatum in the pathophysiology of psychotic disorders (e.g. Howes et al., 2011; Karcher, Rogers & Woodward, 2019) and the limbic striatum in particular is the central region in influential theories of addiction (e.g. Robbins & Everitt, 2002; Everitt & Robbins, 2013). Characterising the effects of THC and CBD on the striatum is therefore vitally important for understanding its role in the pathophysiology of these disorders, and as a means to evaluate potential cannabinoid treatments.

We therefore sought to investigate the effects of cannabinoids on functional connectivity of the striatum, using resting-state fMRI. Firstly, we examined the effects of vaporised herbal cannabis with and without CBD on connectivity in three striatal sub-divisions. In a second study, to isolate the effects of CBD, we investigated the effects of oral CBD vs. placebo in the same regions. Our first hypothesis was that THC will disrupt/reduce striato-cortical functional connectivity particularly in the limbic striatal sub-division. Our second hypothesis was that CBD would ameliorate these effects when delivered in combination with THC. Our third hypothesis was that CBD administered alone would produce a qualitatively different pattern of functional modulations to THC or THC+CBD.

## Methods

### Study 1

Additional data from this study have been published elsewhere (Lawn et al., 2016; Freeman et al., 2017; Wall et al., 2019). These previous reports did not focus on striato-cortical connectivity.

#### Study Design

This study included three drug conditions: cannabis containing both THC and CBD (THC+CBD), high-THC cannabis without CBD (THC) and placebo cannabis (without either THC or CBD). These three conditions were used in a randomized, crossover, placebo-controlled, double-blind design. A Latin Square design was used to randomly assign participants to one of three condition orders. To avoid carry-over effects the scanning sessions were separated by at least 1 week, which is more than three times the elimination half-life of THC (Hindocha et al., 2015).

#### Participants

Seventeen healthy volunteers (9 women) between 18 and 70 years old were recruited (mean age = 26.2, SD = 7.1). The recruitment followed the inclusion criteria for cannabis use of ≤ 3 times per week and ≥ 4 times in the past year. The participants reported on average 8.1 (SD = 5.5) days/month of cannabis use.

Volunteers were excluded if there was current or past history of psychosis in themselves or an immediate family member and if there were any other medical problems considered clinically significant for the study. Additionally, drug related exclusion criteria were previous negative experiences with cannabis, alcohol use was > 5 times per week and use of any other illicit drug > twice per month. For full demographic data, see Lawn et al. (2016). The study was conducted in accordance with the Declaration of Helsinki and was approved by the University College London (UCL) Ethics Committee. Participants provided written informed consent prior to the first study session and they were reimbursed for their time.

#### Drug Administration

All three varieties of cannabis were sourced from Bedrocan (The Netherlands), and were matched for appearance and smell. In each session the same amount of cannabis was administered (133.4 mg). The THC and CBD doses for the current study were determined based on previous experiments that used similar vaporisation methods (Bossong et al., 2009; Hindocha et al., 2015) and Bedrocan product potencies (Niesink et al., 2015). The dose was 8mg THC in both cannabis conditions (THC, THC+CBD) and 10mg of CBD in the THC+CBD condition. The THC (8mg) dose has produced subjective, cognitive, and psychotomimetic effects in previous studies and reflects 1.6 standard units of THC at 5mg (Freeman & Lorenzetti, 2020). All the cannabis was used within 6 months of purchase and was stored in foil-sealed pouches at −20°C and then at ambient temperature immediately prior to administration.

Each cannabis dose was administered using a Volcano Medic Vaporizer (Storz and Bickel, Tuttlingen, Germany) in line with previous studies (Bossong et al., 2009; Hindocha et al., 2015; Mokrysz et al., 2016). The drug was vaporised at 210°C and the product was collected in two balloons. Participants were asked to inhale the drug from the balloons at their own pace and hold each inhalation for 8 seconds.

#### Procedure

Participants completed a telephone screening and then attended a screening visit to assess eligibility, drug history and complete trait questionnaires. In addition they received task training for tasks reported elsewhere (Lawn et al., 2016; Freeman et al., 2017) and a video training of the drug inhalation process. Prior to each study visit, participants were asked to abstain from drug and alcohol use for 24 hours. At the beginning of each visit, a urine test was used to verify the participant’s self-reported drug use and screen for pregnancy. Then the drug was administered and 30 minutes post-administration the MRI scanning session commenced, which lasted approximately one hour. Following the MRI session, participants received a top-up administration and completed a battery of behavioural tasks (reported in Lawn et al., 2016; Mokrysz et al., 2020). Blood samples for measurement of drug concentrations in the plasma were not collected in this experiment.

#### MRI acquisition

A Siemens Avanto 1.5T scanner (Erlangen, Germany) using a 32-channel phased-array head-coil was used to acquire the MRI data. The resting-state functional images were acquired with a T2* gradient-echo echo-planar imaging (EPI) sequence with (TR = 2800 ms, 32 slices, 3.2 mm isotropic voxels, TE = 43 ms, flip-angle = 90°). The scan duration was 12 minutes and 8 seconds, with a total of 260 volumes. At the beginning of the scan session, standard MPRAGE (Magnetization Prepared RApid Gradient Echo) T1-weighted anatomical scans were also acquired for the purposes of co-registration of the functional images (TR = 2730 ms; TE = 3.57 ms; matrix = 176 × 256 × 256; 1 mm isotropic voxels; flip angle = 7°; bandwidth = 190 Hz/pixel; parallel imaging acceleration factor = 2).

### Study 2

Additional data from this study have been published elsewhere; these previous reports did not investigate resting-state striato-cortical connectivity (Bloomfield et al., 2020; Lawn et al., 2020).

#### Study design

The study used a double-blind, randomised, placebo-controlled, repeated-measures design to compare the effects of oral CBD 600mg (pure synthetic (-)-CBD) with matched placebo (PBO) in identical capsules at two sessions. Drug order was completely concealed from participants and experimenters until data collection, entry and analysis had been completed. To avoid carry-over effects the scanning sessions were separated by at least 1 week, which is more than three times the elimination half-life of THC (Hindocha et al., 2014). The order of drug was block randomised and stratified for sex. This study was conducted in accordance with Good Clinical Practice and the Helsinki Declaration (UCL Research Ethics Committee 3325/002). Participants provided written informed consent and received an honorarium for participation (£10 per hour).

#### Drug administration

Synthetic CBD (99.9% purity) was obtained from STI Pharmaceuticals (Brentwood, UK) and manufactured by Nova Laboratories (Leicester, UK). Size 2 gelatin capsules contained microcrystalline cellulose filler and CBD. Matched placebo capsules contained lactose filler. The CBD was formulated in 50 mg capsules. Participants swallowed all 12 capsules at their own pace under invigilation of the experimenter. The 600 mg dose was chosen as it produces an increase in plasma concentrations after acute administration (Englund et al., 2013; Babalonis et al., 2017), is well tolerated in humans (Grotenhermen, Russo & Zuardi, 2017), has been found to produce a significant anxiolytic effect (Bergamaschi et al., 2011b), and has opposing effects to THC on the striatum during fMRI (Bhattacharyya et al., 2010). Previous research suggests that CBD reaches the peak level of plasma concentration after approximately 2.5 hours (Babalonis et al., 2017).

#### Participants

Participants were recruited through online adverts, posters and word-of-mouth. We tested 28 healthy participants. Four participants did not complete both study visits, and one additional subject attended both visits but did not complete the scanning session, so their resting-state data was incomplete. These five subjects were excluded which meant 23 complete sets of data were available for analysis. Subjects ranged in age between 19 and 36 (mean=23.8, SD=4.3), all had normal BMI (mean=22.4, SD=3.6), and had sub-clinical scores on the BDI (mean=2.3, SD=2.9) and BAI (mean=2.6, SD=3.3). No participant showed any evidence of alcohol or nicotine dependence as measured by the AUDIT (mean=1.9, SD=2.1), and the FTND (mean=0, SD=0). All participants included were right-handed and aged 18–70. Exclusion criteria were: (a) current use of psychotropic agents; (b) current or past use of cannabis or CBD; (c) previous use of any psychoactive (recreational) drug on >5 occasions; (d) current or previous mood disorder, psychosis, anxiety disorder, or substance abuse disorder according to Diagnostic and Statistical Manual of Mental Disorders IV (DSM-IV) criteria; (e) current nicotine dependence (defined by Fagerström Test for Nicotine Dependence; Heatherton, Kozlowski & Fagerström, 1991); (f) score >7 on the Alcohol Use Disorders Identification Test (Saunders et al., 1993); (g) pregnancy; (h) impaired mental capacity; (i) allergy to CBD or placebo excipients; (j) claustrophobia or other contraindications to MRI.

#### Procedure

Participants completed a screening on the telephone during which initial eligibility criteria (drug use, FTND, AUDIT, MRI contraindications, allergies, medical information and handedness) were assessed and basic participant details were recorded. Participants who appeared eligible on the phone were invited to attend experimental sessions. Participants were asked to fast from midnight the day before both sessions, and refrain from smoking tobacco and consuming alcohol for 24 h before the start of the sessions. Upon arrival, participants underwent urine tests to verify they were not pregnant (if female) and they had not recently taken recreational drugs. They also completed breath tests for alcohol and carbon monoxide. Eligible participants then completed two seven-hour experimental sessions, when they received CBD or placebo on the first session, and the other drug condition on the second session. The MRI scanning session commenced 2.5 hours after drug administration and lasted approximately 1.5 hours.

#### Plasma CBD concentrations

We performed venipuncture immediately after MRI scanning to measure CBD concentrations. Blood samples were collected in EDTA vacutainers and were immediately centrifuged to plasma for storage at −80°C. Samples were analysed using Gas Chromatography coupled with Mass Spectrometry with a lower limit of quantification of 0.5 ng/mL.

#### MRI acquisition

MRI data was collected using a 3-Tesla Siemens Prisma MRI Scanner at the Robert Steiner MR unit at Hammersmith Hospital, London. Functional imaging used a multiband (acceleration factor= 2) gradient-echo T2*-weighted echo-planar imaging (EPI) sequence with 42 slices per volume (Repetition time [TR]=2400 ms; Time to Echo [TE]=30 ms; in-plane matrix=64×64; 3 mm isotropic voxels; flip angle=62°; bandwidth=1594 Hz/pixel; 304 volumes; a slice thickness of 3 mm; field of view=192 × 192 mm). The phase encoding direction was from anterior to posterior. There were three dummy scans at the beginning of the scan, which were not included in our dataset. For structural acquisition, a T1-weighted structural volume was acquired for all participants using a MPRAGE scan (TR=2300 ms; TE=2.28 ms, TI=900 ms, flip angle=9°, field of view= 256 mm, image matrix=256 with 1 mm isotropic voxels; bandwidth=200 Hz/pixel).

### Statistical analysis (Study 1 and 2)

Image analyses were performed using FSL 5.0.4. The functional data were pre-processed using spatial smoothing with a 6 mm FWHM (full-width, half-maximum) Gaussian kernel, high-pass temporal filtering (100 s), head motion correction using MCFLIRT and non-linear registration to a standard template (MNI152). The anatomical data were skull-stripped using FSL’s brain extraction tool (BET) and segmented using FMRIB’s automated segmentation tool (FAST), into grey/white matter and cerebro-spinal fluid (CSF).

#### Striatal Networks: Seed-based analysis (Study 1 and 2)

Brain masks for the three striatal networks (associative, limbic and sensorimotor) were defined according to the original definition by Martinez et al., (2003), and using the atlas provided by (Tziortzi et al., 2013). The associative mask included the precommissural dorsal caudate, the precommissural dorsal putamen and postcommissural caudate. The limbic mask included the ventral pallidum and substantia nigra; and the sensorimotor mask comprised the postcommissural putamen.

A set of seed-based analyses were conducted using methods similar to previous reports (Demetriou et al., 2016; Comninos et al., 2018; Wall et al., 2019). The standard-space striatal brain masks were co-registered to each participant’s functional image space, and time-series were extracted from these regions that were subsequently used in the first-level analysis models as regressors of interest. Additionally, the white matter and CSF time-series from each participant were included in the analysis models as regressors of no interest, along with head-motion regressors. First-level models included use of FSL’s FILM algorithm to correct for auto-correlation in the time-series. Higher-level analyses were performed using FSL’s FLAME-1 mixed-effects model, and results were cluster-corrected for multiple comparisons at *Z* >2. 3, *p* <.05. Separate group-level models were produced in order to model mean functional connectivity effects (all subjects, all scans) for each study, and voxelwise comparisons between the drug conditions (three comparisons in study 1, two in study 2). To quantify the condition effects across each striatal network, the group mean functional connectivity results were used to produce image masks (thresholded at *Z*=5) from which numeric data were extracted for each subject/scan. Drug effects on mean network connectivity were assessed using 2-tailed paired *t* tests with a corrected alpha level of *p* < 0.008 in order to account for multiple comparisons.

## Results

### Study 1

#### Seed-based functional connectivity analyses

There were no effects seen in the active drug conditions > placebo contrasts, in any of the analyses, meaning the conditions did not significantly increase connectivity relative to placebo. When administered alone, THC significantly disrupted (placebo > active conditions) mean connectivity between the limbic striatum and the bilateral insula and frontal opercular cortex as shown in Figure 1. By contrast, when THC was co-administered with CBD there was no evidence for disruption of connectivity between limbic striatum and any brain region.

**Figure 1.**
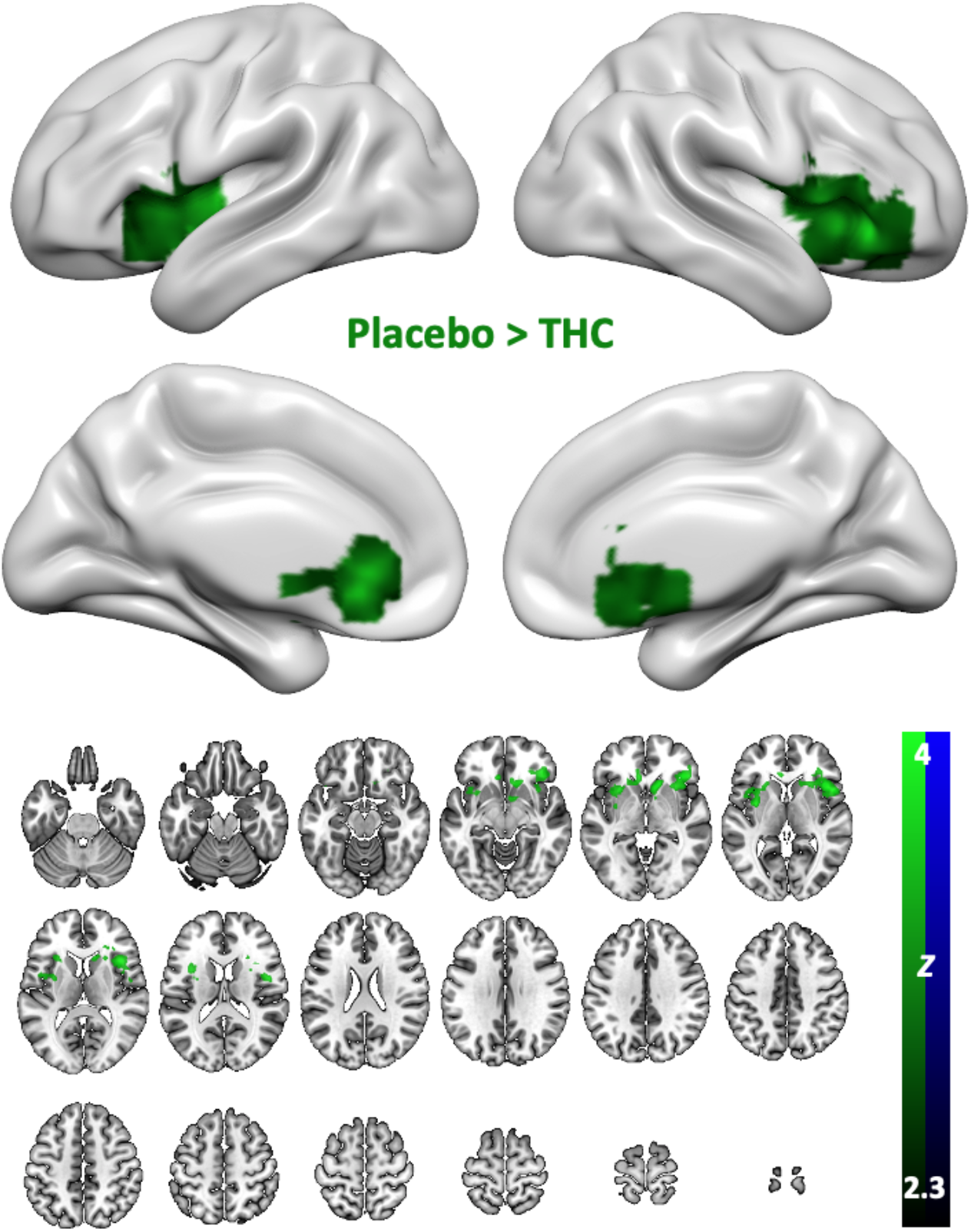
Drug effects on brain wide connectivity with the limbic striatum in study 1. Contrast is placebo > THC. Clusters represent a decrease in functional connectivity with the limbic striatum in the active drug condition. The THC+CBD condition showed no significant effects for this seed-region.

Administration of the THC+CBD condition reduced connectivity of the associative striatum with the dorsal anterior cingulate as well as a large lateral region covering part of frontal opercular cortex and sensorimotor regions in the left hemisphere (more restricted in the right hemisphere). The THC condition showed a broadly similar, though somewhat more widespread) distribution, with the regions affected covering more of the frontal operculum and extending downwards into the insula. See Figure 2.

**Figure 2.**
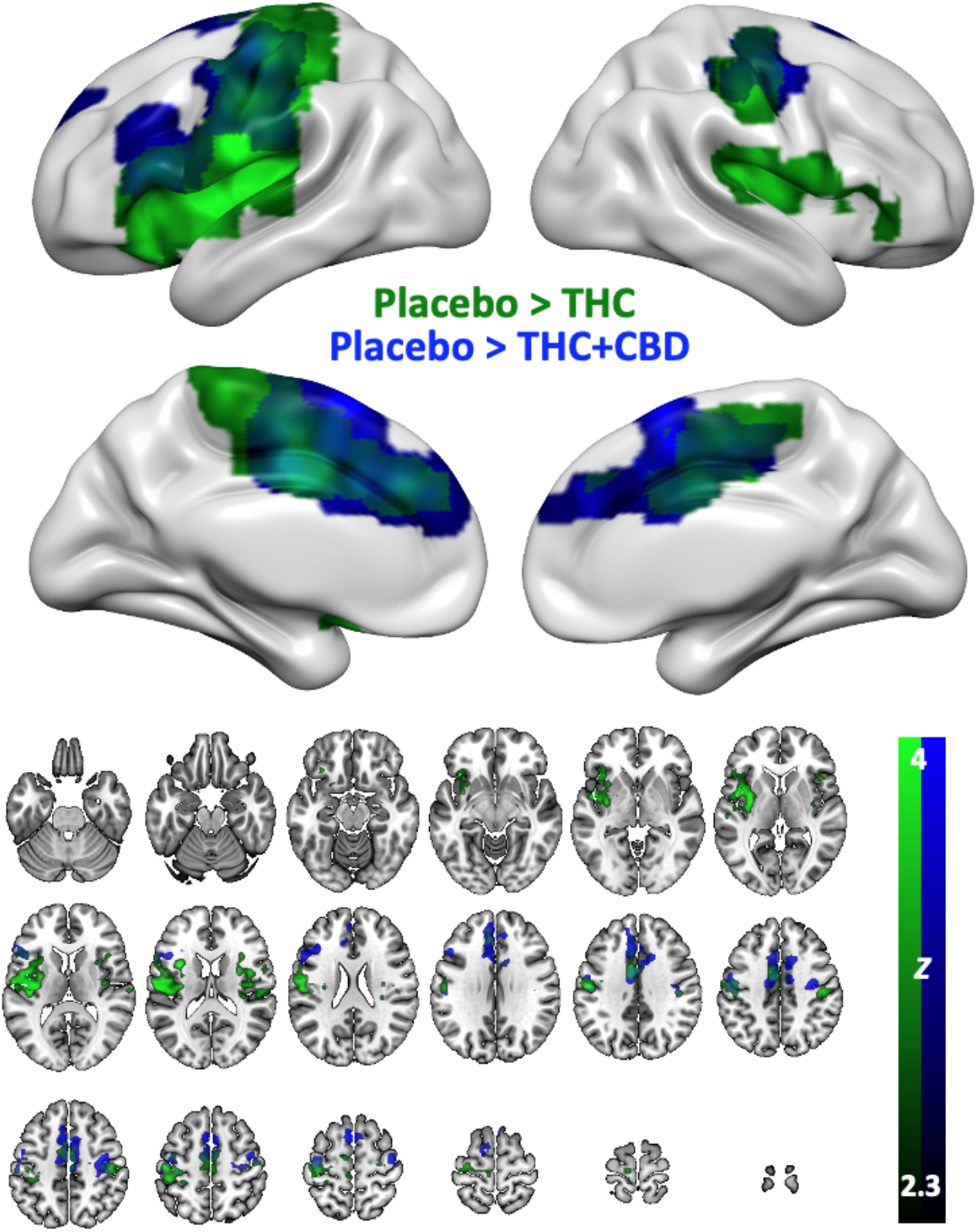
Drug effects on brain wide connectivity with the associative striatum in study 1. Contrasts are placebo > active drug. Clusters represent a decrease in functional connectivity with the associative striatum in the active drug conditions. The green scale shows the THC condition and the blue scale shows THC+CBD.

Connectivity with the sensorimotor striatum was the most strongly disrupted of the striatal networks in this study. The THC+CBD condition reduced activity within many sensory-motor associated areas such as the parietal operculum cortex, central opercular cortex and the post central gyrus. Language and auditory associated areas also had reduced connectivity including the supramarginal gyrus, planum temporale and Heshcl’s gyrus. There was also some reduction seen in the motor cortex. Similar disruptions were seen in the THC condition, the most notable differences are larger portion of Heschl’s gyrus disrupted as well as secondary somatosensory cortex.

**Figure 3.**
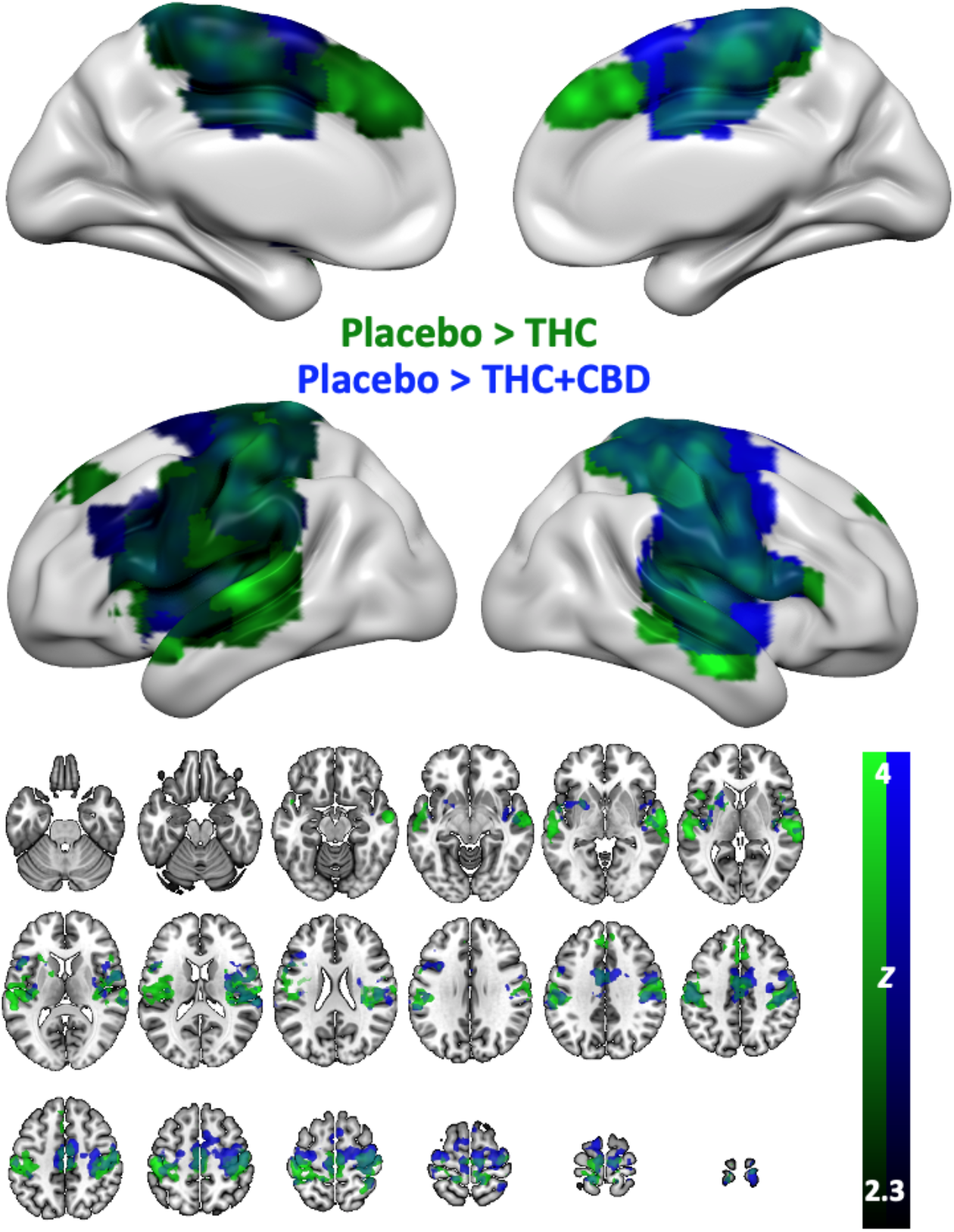
Drug effects on brain wide connectivity with the sensorimotor striatum in study 1. Contrasts are placebo > active drug. Clusters represent a decrease in functional connectivity with the sensorimotor striatum in the active drug conditions. The green scale shows the THC condition and the blue scale shows THC+CBD.

The overall mean connectivity of each network was also examined using thresholded versions of the group-mean connectivity maps as mask images. The largest effect of the active conditions (relative to placebo) was in the sensorimotor network (THC+CBD: *t*[16] = 2.93, *p* = .01; THC: *t*[16] = 3.07, *p* = .007).

### Study 2

Results from Study 2 showed a markedly different effect of oral CBD on striatal functional connectivity. Figure 5 shows results from all three analyses (using associative, limbic, and sensorimotor subdivisions as seed regions) and for the CBD condition vs. placebo. Connectivity analyses with the associative sub-division showed drug effects in bilateral areas in the posterior parietal lobes, extending medially into the parieto-occipital sulcus and into the posterior cingulate in the left hemisphere. It is important to note that this result is the opposite contrast to the results found in study 1 (and in fact, the other two results described below from study 2), and is in fact CBD > placebo, implying a relative *increase* in functional connectivity between these regions and the associative striatum, under the CBD condition. No areas showing significant relative decreases (placebo > CBD) were found in this analysis. For the limbic striatum seed-region, an area of decreased connectivity (placebo > CBD) was found in the right hemisphere insula and lateral frontal cortex. For the sensorimotor seed region, significant clusters of relatively decreased connectivity (placebo > CBD) were seen in the left cerebellum. For these latter two analyses, no areas showing significant relative increases (CBD > placebo) were found.

**Figure 5.**
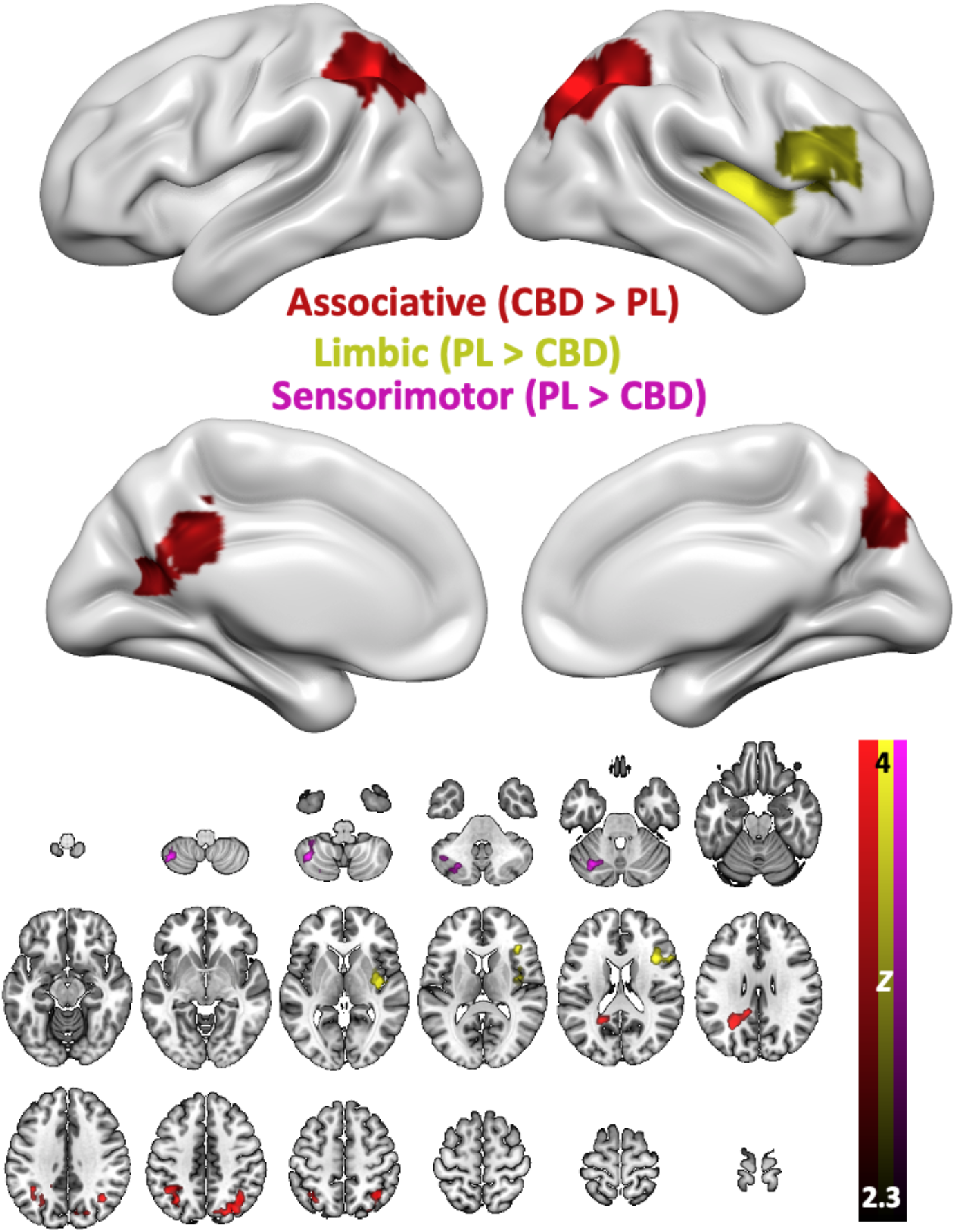
Drug effects on brain wide connectivity with the associative (red), limbic (yellow), and sensorimotor (pink) striatum in study 2. Both relative increases (CBD > PL) and decreases (PL > CBD) are shown, depending on the pattern of significant results in the three analyses. Effects on sensorimotor striatum connectivity were only seen in the cerebellum, and are therefore not visible on the top panel, which only shows inflated views of the cortex.

## Discussion

The present data demonstrate extensive effects of cannabinoids on striatal functional connectivity networks. In study 1, effects on the limbic striatum were specific to the THC condition, with disruptions (relative decreases in connectivity with the active drug condition) seen in the anterior insula, and areas of the striatum itself. Effects of the different drug conditions on associative striatal connectivity were both widespread, and somewhat dissociated, with both strains disrupting dorsal regions (ACC and motor cortex) but the THC condition also specifically affecting more ventral regions (frontal operculum and insula). Regions affected in the sensorimotor striatum analysis were somewhat similar, with perhaps less of a dorsal/ventral dissociation between the two conditions. In study 2, the effect of 600mg CBD is noticeably weaker and less widespread, with disruption of connectivity in the analyses of limbic and sensorimotor seed-regions only seen in localised regions in one hemisphere (the insula/lateral frontal lobe, and the cerebellum, respectively). Intriguingly, the analysis of the associative striatum connectivity in study 2 showed a result of opposite polarity; a relative increase, or enhancement of connectivity, in parietal regions as a result of the drug administration.

Overall, it is clear cannabinoids can have profoundly disruptive effects on striatal functional connectivity, but the effects of CBD alone are relatively minor, and the effects of THC are effectively blocked by the presence of CBD in the limbic striatum. Even in the associative and sensorimotor striatum, effects of the THC-only condition (THC) in study 1 are more widespread, also suggesting that CBD is moderating the effect of THC in these networks to some extent. The finding in study 2 that CBD actually increases associative striatum connectivity is consistent with the result in study 1 of an ameliorating effect of the CBD on the disruptive effects of THC in the associative striatum, when administered together. The specific effect of the pure-THC (THC) condition on the limbic striatum here is mirrored by a key previous result (Bossong et al., 2015) which showed that only the limbic striatum showed reliable dopamine release with a THC challenge, indexed by [^11^C]raclopride Positron Emission Tomography (PET). This study used synthetic (therefore, pure) THC as the acute challenge; the present data therefore extend this result by suggesting that CBD may potentially block the release of dopamine produced by THC in the limbic striatum. CBD alone may also have effects on limbic striatum connectivity, as seen in study 2, where the (right) insula is also significantly modulated by the oral CBD condition.

This may be significant, as the limbic striatum consists of the nucleus accumbens and the head of the caudate. The nucleus accumbens is one of the primary substrates known to be heavily involved in the formation and maintenance of addiction (Robinson & Berridge, 1993, 2001; Robbins & Everitt, 2002; Volkow et al., 2007). The increasing concentration of THC in modern cannabis (which also often has relatively low-levels of CBD; Niesink et al., 2015; El Sohly et al., 2016) is thought to be a major factor in the increase of cannabis related-health issues, in particular addiction (Freeman & Winstock, 2015). The finding here that CBD blocks the disruptive effect on limbic striatum connectivity is also consistent with previous behavioural work showing that CBD attenuates the appetitive and incentive-salience effects of THC and other drugs (Morgan et al., 2010; Hindocha et al., 2018). Taken together these various findings suggest a possible physiological mechanism whereby THC promotes dopamine release in the ventral striatum, making users who consume relatively pure THC strains vulnerable to addiction. However, in users of more balanced strains containing CBD, the acute dopaminergic and addiction-promoting effects of THC on the ventral striatum are ameliorated, or perhaps blocked entirely. This ‘buffering’ effect of CBD is also consistent with the previous results reported from this cohort (Lawn et al., 2016; Freeman et al., 2017; Wall et al., 2019).

The finding of a relative increase in connectivity with the CBD condition (in the associative striatum analysis) is mirrored by a recent similar finding in Grimm et al. (2018), which also used oral administration and the same dose as the present data (600mg). These authors showed a relative increase in frontal-striatal connectivity with CBD, and speculate that this might account for the anti-psychotic effects of CBD, as fronto-striatal connectivity effects are a common finding in studies of schizophrenic patients (e.g. Fornito et al., 2013). Another converging result is that of Rzepa, Tudge & McCabe (2016) which used the CB1 neutral antagonist tetrahydrocannabivarin (THCV). This study showed increased connectivity within the executive control network; usually conceived as a network subserving attentional and cognitive processes involved in task engagement. Cannabidiol also may be a negative allosteric modulator at CB1 receptors (Laprairie et al., 2015; Chesney et al., 2020), and here we show increases in connectivity in the associative striatum; the region of the striatum most associated with cognitive functions and brain regions.

We also see marked effects on the insula, across all three networks examined in study 1, and for the limbic striatum network in study 2. The insula is a key hub in the salience network (Seeley et al., 2007; Goulden et al., 2014; Uddin, 2014) and recent work using combined PET and fMRI methods has identified a link between mesolimbic dopamine systems and the salience network (McCutcheon et al., 2019b). Connectivity between the striatum and the salience network has also been shown to be affected in psychotic disorders (Karcher, Rogers & Woodward, 2019), and striatal-salience network connectivity has been shown to be increased in individuals exposed to chronic psychosocial stressors (a key hypothesised factor in the development of psychosis; McCutcheon et al., 2019a). Taken together, these findings suggest a clear role for striatal-salience network connectivity in the pathophysiology of psychotic disorders, and further suggest that compounds that specifically target these systems (such as CBD) may be useful therapeutically.

To the authors’ knowledge, this is the first report in human subjects of a comparison of THC, THC+CBD and CBD, achieved using a unified set of analysis methods, and with all comparisons performed in a placebo-controlled, double-blind design. These are important strengths, however, as the data come from two separate studies a direct comparison between each of the conditions is compromised by the use of different cohorts of subjects, and different routes of administration (inhalation in study 1, oral dosing in study 2) and doses. Other differences between studies were scanner model and field strength (1.5 Tesla in study 1, 3 Tesla in study 2), data acquisition protocol, and length of the scan.

## Conclusion

Cannabinoids exert a major acute effect on striato-cortical functional connectivity, with effects on striatal connectivity with the insula particularly evident across all three drug conditions. These effects on the limbic striatum in particular and its connectivity with the insula (and by implication, the salience network) are likely a crucial finding in our evolving understanding of the acute brain effects of cannabinoids. A key question for future research is understanding how these acute effects translate into longer-term effects in chronic users, what role these striato-cortical connections may have in the pathophysiology of cannabis use disorder and cannabis-related psychosis, and what therapeutic options might usefully target them. These questions will grow increasingly more urgent as cannabis seems likely to continue its transition to quasi-legal or fully-legal status in a growing number of jurisdictions.

## Supplementary Figures

**Figure S1.**
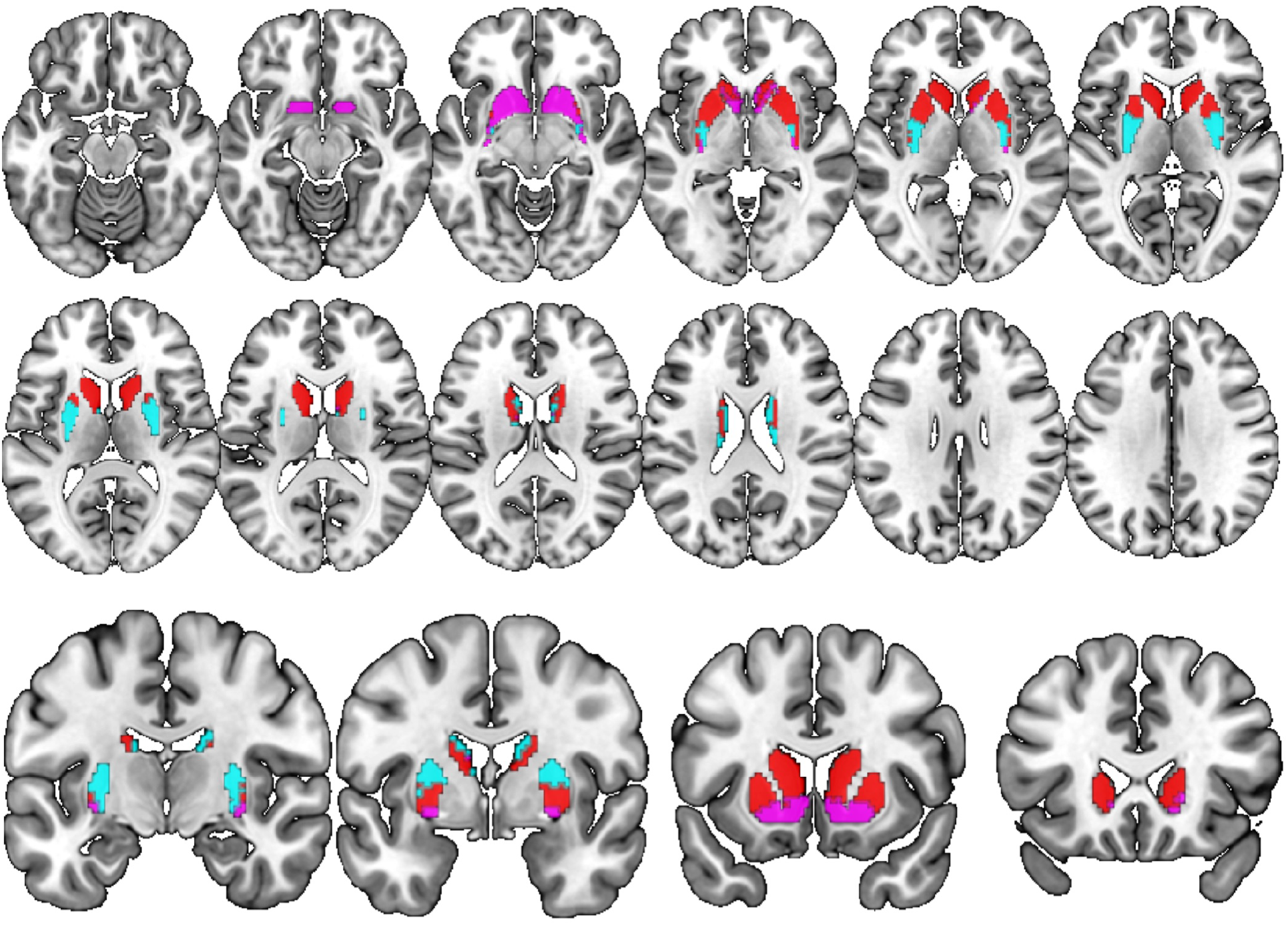
Seed regions used in the functional connectivity analyses, derived from the atlas provided by Tziortzi et al. (2013). Associative striatum in red, limbic striatum in pink, and sensorimotor striatum in cyan.Associative network

**Figure S2.**
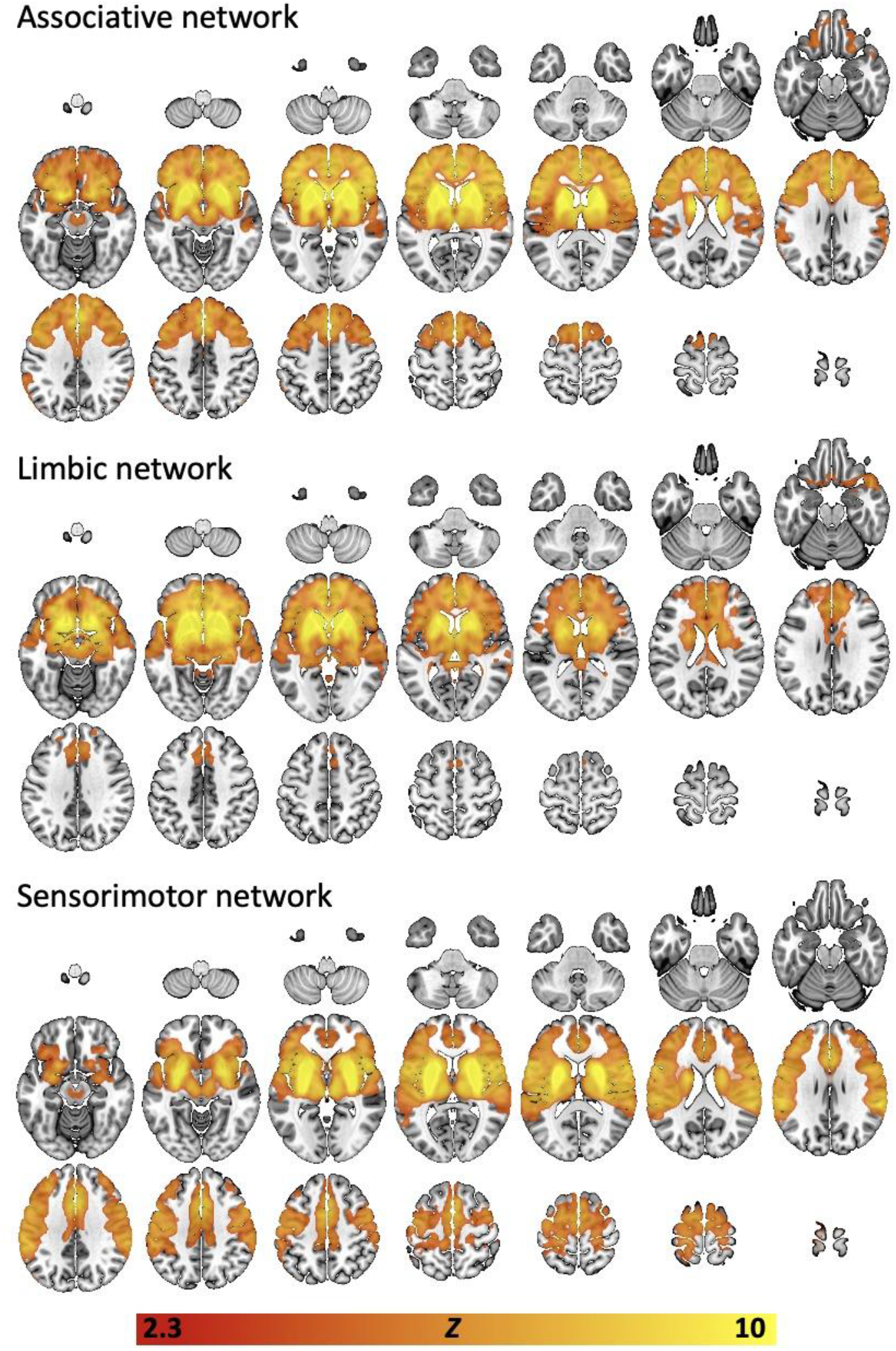
Group-mean (all subjects, all scans) connectivity networks derived using the seed-regions shown in figure S1, and the resting-state fMRI data from study 1 (N=17). Top panel = associative network, middle panel = limbic network, bottom panel = sensorimotor network. Statistical thresholds are *Z*=2.3, *p* < 0.05 (cluster-corrected).

**Figure S3.**
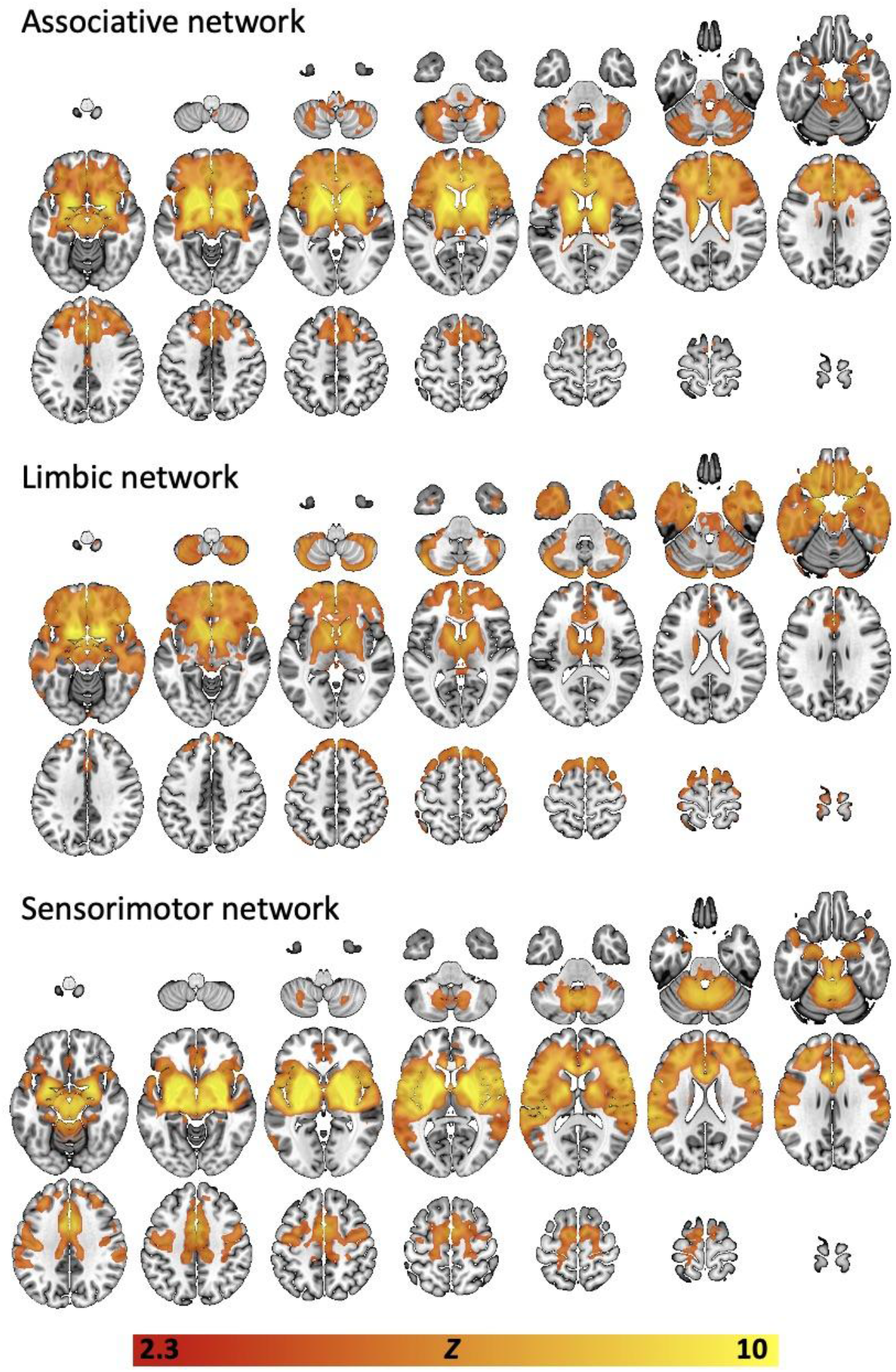
Group-mean (all subjects, all scans) connectivity networks derived using the seed-regions shown in figure S1, and the resting-state fMRI data from study 2 (N=23). Top panel = associative network, middle panel = limbic network, bottom panel = sensorimotor network. Statistical thresholds are *Z*=2.3, *p* < 0.05 (cluster-corrected).

